# Epigenetic changes in autoimmune monocytes contribute to disease and can be targeted by JAK inhibition

**DOI:** 10.1101/2021.11.20.469222

**Authors:** J.G.C. Peeters, A. Boltjes, R.C. Scholman, S.J. Vervoort, P.J. Coffer, M. Mokry, S.J. Vastert, F. van Wijk, J. van Loosdregt

## Abstract

How the local inflammatory environment regulates epigenetic changes in the context of autoimmune diseases remains unclear. Here we assessed the transcriptional and active enhancer profile of monocytes derived from the inflamed joints of Juvenile Idiopathic Arthritis (JIA) patients, a model well-suited for studying autoimmune diseases. RNA-sequencing analysis demonstrated that synovial-derived monocytes display an activated phenotype, characterized by increased expression of surface activation markers and multiple chemokines and cytokines. This increased expression is regulated on the epigenetic level, as ChIP-sequencing indicated increased H3K27 acetylation of genes upregulated in synovial monocytes. IFN signaling-associated genes are increased and epigenetically altered in synovial monocytes, suggesting a role for IFN in establishing the inflammatory phenotype. Treatment of human synovial monocytes with the Janus-associated kinase (JAK) inhibitor Ruxolitinib, which inhibits IFN signaling, transformed the activated enhancer landscape and reduced disease-associated gene expression, thereby inhibiting the inflammatory phenotype. Taken together, these data provide novel insight into epigenetic regulation of autoimmune disease-patient derived monocytes and suggest that altering the epigenetic profile can be a therapeutic approach for the treatment of autoimmune diseases.

## Introduction

Autoimmune diseases are characterized by abnormal immune responses due to loss of immunological tolerance. The presence of autoreactive T and B cells and autoantibodies often links the pathogenesis of autoimmune diseases to the adaptive immune system, but innate immune cells, such as monocytes, also play a key role. Monocytes can recruit adaptive immune cells to the site of inflammation by secreting chemokines and further perpetuate immune activation, through either direct cell-cell contact or abundant secretion of pro-inflammatory cytokines (i.e. IL-6, TNF-α). As such, many autoimmune diseases are associated with increased monocyte infiltration (*1*–*5*). The exact mechanisms that regulate monocyte activity remain to be determined, but it is clear that environmental factors and epigenetic regulation are important in autoimmune disease pathogenesis (*6*). Enhancers are *cis*-regulatory DNA regions, generally a few hundred base pairs in size, that are pivotal for the spatio-temporal regulation of gene expression by recruiting RNA polymerase II, transcription factors and co-factors, such as the histone acetyltransferase p300 or the mediator complex (*7*). Enhancers regions are characterized by methylation of histone H3 at lysine 4 (H3K4me) and contain H3K27 acetylation in their active status (*8, 9*). Around 60% of the single nucleotide polymorphisms (SNPs) associated with autoimmune disease are localized in enhancer regions and disease-associated variants preferentially map to enhancer regions specific for disease-relevant cell types (*10*–*15*). For example, SNPs associated with multiple sclerosis (MS) and systemic sclerosis (Ssc) have been demonstrated to map to monocyte enhancer regions, indicating that epigenetic regulation plays an important role in autoimmune disease pathogenesis (*16*). In addition, alterations of the histone modification profile have been described in monocytes and other autoimmune disease patient-derived cells (*17*–*19*).

Information concerning epigenetic changes specifically at the site of inflammation is currently lacking. Reasons for this include the difficulty in obtaining enough immune cells from the local inflammatory site to perform in-depth analyses. For regulatory and effector T cells it has been demonstrated that the epigenetic landscape is altered at the site of autoinflammation and that these changes contribute to their pro-inflammatory phenotype (*12, 20*). These studies indicate that the inflammatory environment can directly induce epigenetic changes that result in a pro-inflammatory phenotype that contributes to autoimmune pathogenesis. The role of epigenetic regulation in monocytes at the site of inflammation has not been well studied. Here, we evaluated role of active enhancers in monocytes in a human autoimmune-associated inflammatory environment, by using monocytes derived from the inflamed joints of Juvenile Idiopathic Arthritis (JIA) patients. RNA-sequencing and chromatin immunoprecipitation (ChIP) sequencing for H3K27ac demonstrated that human monocytes from the local inflammatory site display an epigenetically-regulated activated phenotype. This phenotype was characterized by increased expression of IFN-responsive genes. Treatment of inflammatory site-derived monocytes with the JAK-1,2 inhibitor Ruxolitinib reduced the active state of various enhancer regions, correlating with impaired expression of disease-associated genes. Taken together, these data demonstrate that IFN signaling induces epigenetic alterations and that these changes contribute to a pro-inflammatory phenotype, suggesting that modulating the epigenetic landscape might be a potential therapeutic approach for the treatment of autoimmune diseases.

## Results

### Synovial-derived monocytes display an activated phenotype on the transcriptional level

To evaluate the transcriptional changes at the site of local autoimmune inflammation, monocytes were collected from inflamed joints of Juvenile Idiopathic Arthritis (JIA) patients. RNA-sequencing analysis of JIA CD14^+^CD16^+^ monocytes obtained from the synovial compartment (SF) and HC peripheral blood (PB)-derived CD14^+^CD16^+^ cells was performed. 2461 genes were significantly differentially expressed (Figure 1A and Supplemental Figure 1A). Genes that were upregulated in monocytes derived from the local inflammatory site are associated with regulation of the immune system and in particular with cytokine responses (Figure 1B and 1C). Furthermore, the top 3 diseases associated with these genes are all autoimmune diseases (Figure 1D). Analysis of monocyte surface markers revealed increased expression of several surface activation markers, including *CD80, CD69, CCR2*, and *FCGR1A* in monocytes (Figure 1E). In addition, many cytokines and chemokines were expressed at a higher level by monocytes derived from the inflammatory environment (Figure 1F). These observations were further validated on both the mRNA and protein level for several cytokines, chemokines and surface markers (Figure 1G, 1H and Supplemental Figure 1B). Collectively, these data demonstrate that transcriptional changes in monocytes result in an activated phenotype associated with (auto-)inflammation.

**Figure 1:**
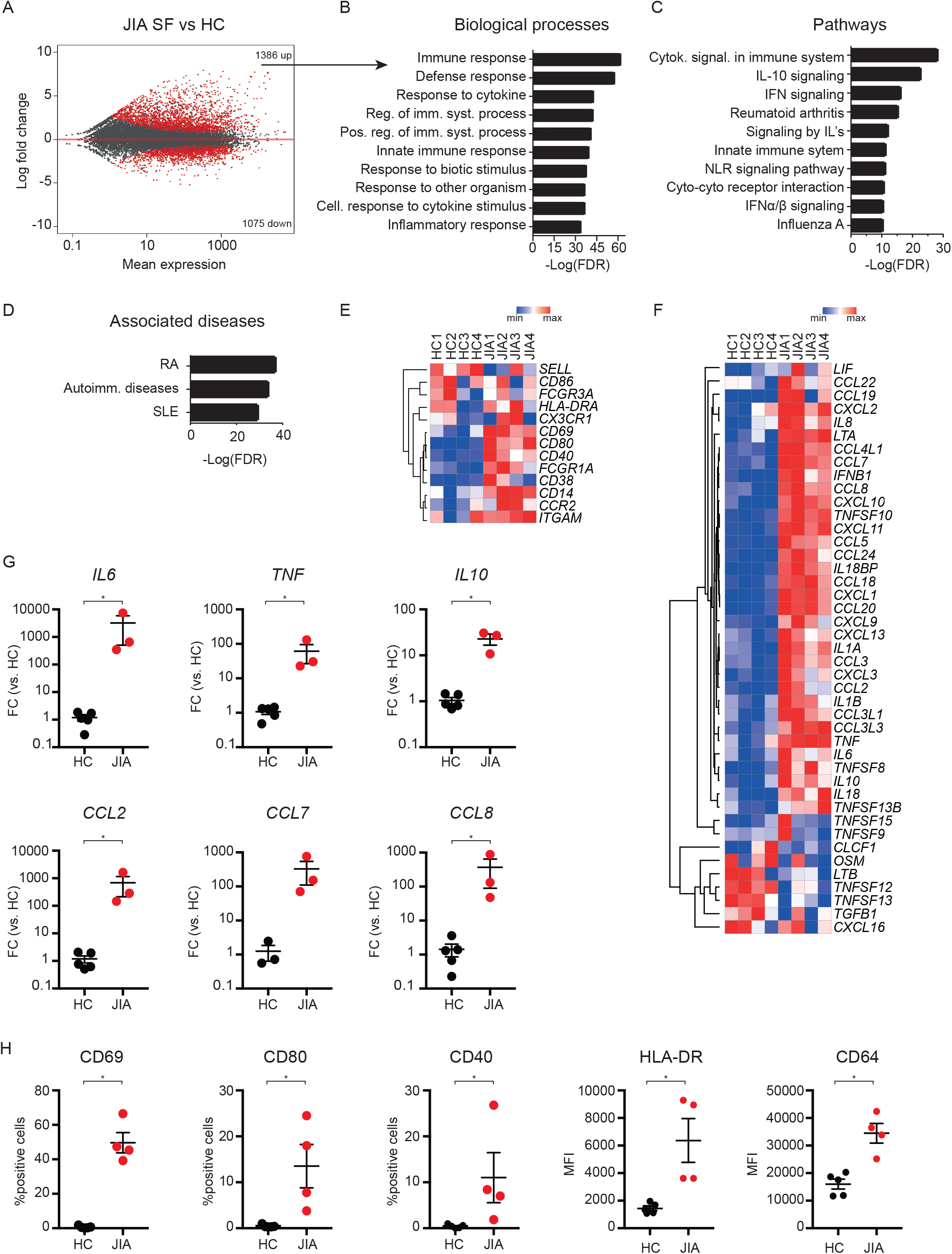
Transcriptional analysis of JIA SF monocytes indicates an activated phenotype. (**A**) MA plot displaying genes differentially expressed between HC PB monocytes and JIA SF monocytes. Red dot indicate genes with a FDR<0.05. (**B**) Top 10 biological processes associated with genes that are significantly increased in JIA SF monocytes. (**C**) Top 10 pathways associated with genes that are significantly increased in JIA SF monocytes. (**D**) Top 3 diseases associated with genes that are significantly increased in JIA SF monocytes. (**E, F**) Heatmap demonstrating the expression of selected monocyte surface markers (**E**) and selected cytokines (**F**) and chemokines in HC PB and JIA SF monocytes. (**G**) Gene expression of selected cytokine and chemokines in JIA SF monocytes vs. HC PB monocytes. (**H**) Protein levels of selected surface markers in HC PB and JIA SF monocytes. P values for F and G were calculated using a Mann-Whitney test. * = p<0.05; ** = p<0.01, *** = p<0.001.

### H3K27ac regulation contributes to the activated phenotype of synovial monocytes

To determine whether the transcriptional changes in synovial monocytes are regulated on the epigenetic level, ChIP-sequencing for H3K27ac, was performed on HC PB-derived monocytes, JIA PB-derived monocytes, and JIA SF monocytes. Principal component analysis demonstrated that JIA SF monocytes clustered separately from HC and paired JIA PB-derived monocytes (PC1: 44%), indicating that their H3K27ac profile is distinct (Figure 2A). Indeed, more than 3000 regions contain a significantly different H3K27ac signal in inflammation-derived monocytes compared to either HC or JIA PB monocytes, while JIA PB and HC PB monocytes are relatively similar regarding their H3K27ac profile (Figure 2B, 2C and Supplemental Figure 2A). Analysis of the genes associated with an increased H3K27ac signal in local inflammatory site-derived monocytes compared to JIA PB monocytes, indicated association with biological processes that are involved in regulation of the immune system and include many cytokine and chemokines of which the expression was upregulated in inflammatory monocytes (Figure 2D-2E). Indeed, gene set enrichment analysis demonstrated that genes associated with an increased H3K27ac signal are enriched within the mRNAs upregulated at the site of inflammation, indicating that an increased H3K27ac signal correlates with increased gene expression in synovial monocytes (Figure 2F). In agreement with this, genes associated with an increased H3K27ac signal in JIA are more abundantly expressed in synovial monocytes compared to HC monocytes (Figure 2G and 2H). To further assess whether changes in H3K27ac contribute to JIA gene expression, the bromodomain and extra-terminal domain (BET) inhibitor JQ1 was used to inhibit enhancer-mediated transcription (*21*). Treatment of synovial monocytes with JQ1 preferentially reduced the expression of genes that were upregulated in JIA monocytes, indicating that expression of these genes depends on H3K27ac regulation (Figure 2I, 2J and 2K and Supplemental Figure 2B and 2C). Altogether, these data indicate that disease-associated gene expression in monocytes derived from the local inflammatory site is mediated by increased H3K27 acetylation.

**Figure 2:**
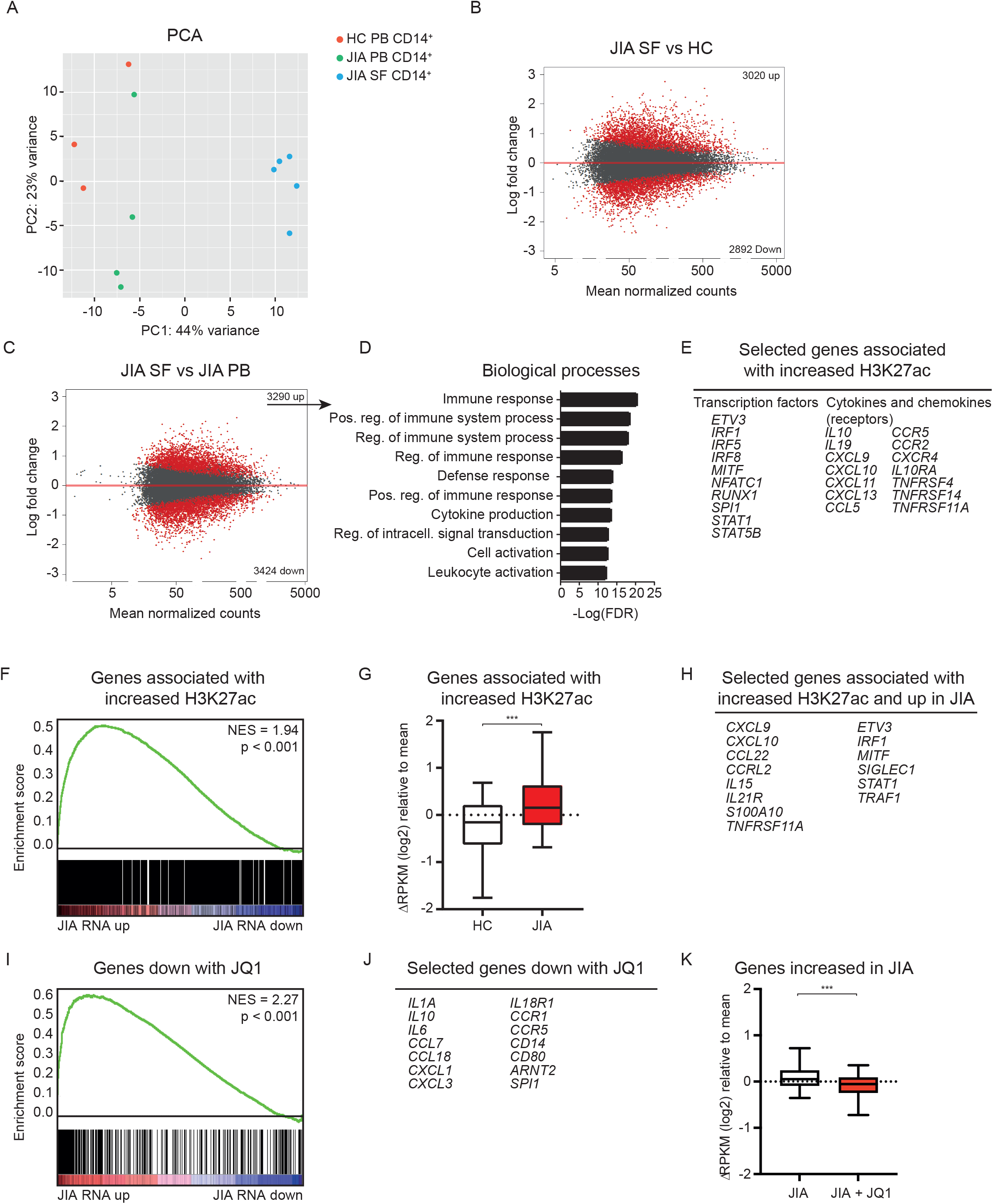
Epigenetic regulation contributes to the activated phenotype of JIA SF monocytes. (**A**) Principal component analysis of H3K27ac signal in HC PB, JIA PB and JIA SF monocytes. (**B and C**) MA plot of H3K27ac regions different between JIA SF and HC PB (left) and between JIA SF and JIA PB monocytes. Red dot indicate H3K27ac regions with a FDR<0.1. (**D**) Top 10 biological processes associated with genes that are significantly increased in JIA SF vs. JIA PB monocytes. (**E**) Selected genes associated with increased H3K27ac signal in JIA SF vs. JIA PB monocytes. (**F**) Gene set enrichment analysis of genes associated with an increased H3K27ac signal in JIA SF monocytes and genes differentially expressed within JIA SF monocytes. (**G**) Boxplot with 5%-95% whiskers displaying ΔRPKM (log2) values of genes associated with an increased H3K27ac signal in HC and JIA monocytes. (**H**) Selected genes associated with an increased H3K27ac signal and increased in JIA. (**I**) Gene set enrichment analysis of genes decreased with JQ1 and genes differentially expressed within JIA SF monocytes. (**J**) Selected genes decreased with JQ1. (**K**) Boxplot with 5%-95% whiskers displaying ΔRPKM (log2) values of genes increased in JIA in JIA samples treated with or without JQ1. P values for G and K were calculated using a Wilcoxon-matched pairs signed rank test. * = p<0.05; ** = p<0.01, *** = p<0.001.

### JAK inhibition alters the H3K27ac profile of synovial monocytes

We observed that genes upregulated in inflammatory monocytes are associated with IFN signaling (Figure 1C). Detailed analysis of IFNγ signaling-associated genes (based on REACTOME IFN signaling pathway) confirmed that a majority of these genes are indeed upregulated in monocytes derived from the inflammatory environment (Figure 3A). RT-qPCR analysis validated the increased expression of several IFN-induced genes and the same trend was observed on the protein level (Figure 3B and Supplemental Figure 3A). Pathway analysis of the genes associated with an increased H3K27ac signal in inflammatory monocytes indicated an association with IFNγ signaling (Figure 3C). IFN signaling is mediated by the Janus Kinase (JAK) and Signal Transducer and Activator of Transcription (STAT) pathway (*22*). Gene set enrichment analysis demonstrated that genes belonging to the JAK-STAT signaling pathway are upregulated in synovial monocytes, suggesting that this pathway may regulate gene expression at the site of inflammation (Figure 3D). Similarly, we observed increased expression of STAT1 and STAT2 and increased H3K27 acetylation of a STAT1 and STAT2-associated enhancer (Figure 3E and 3F). To further define the role of the JAK-STAT signaling pathway, we assessed the H3K27ac and STAT1 ChIP-sequencing dataset of IFNγ-treated human primary monocytes generated by Qiao *et al* (GSE43036, (*23*)).

**Figure 3:**
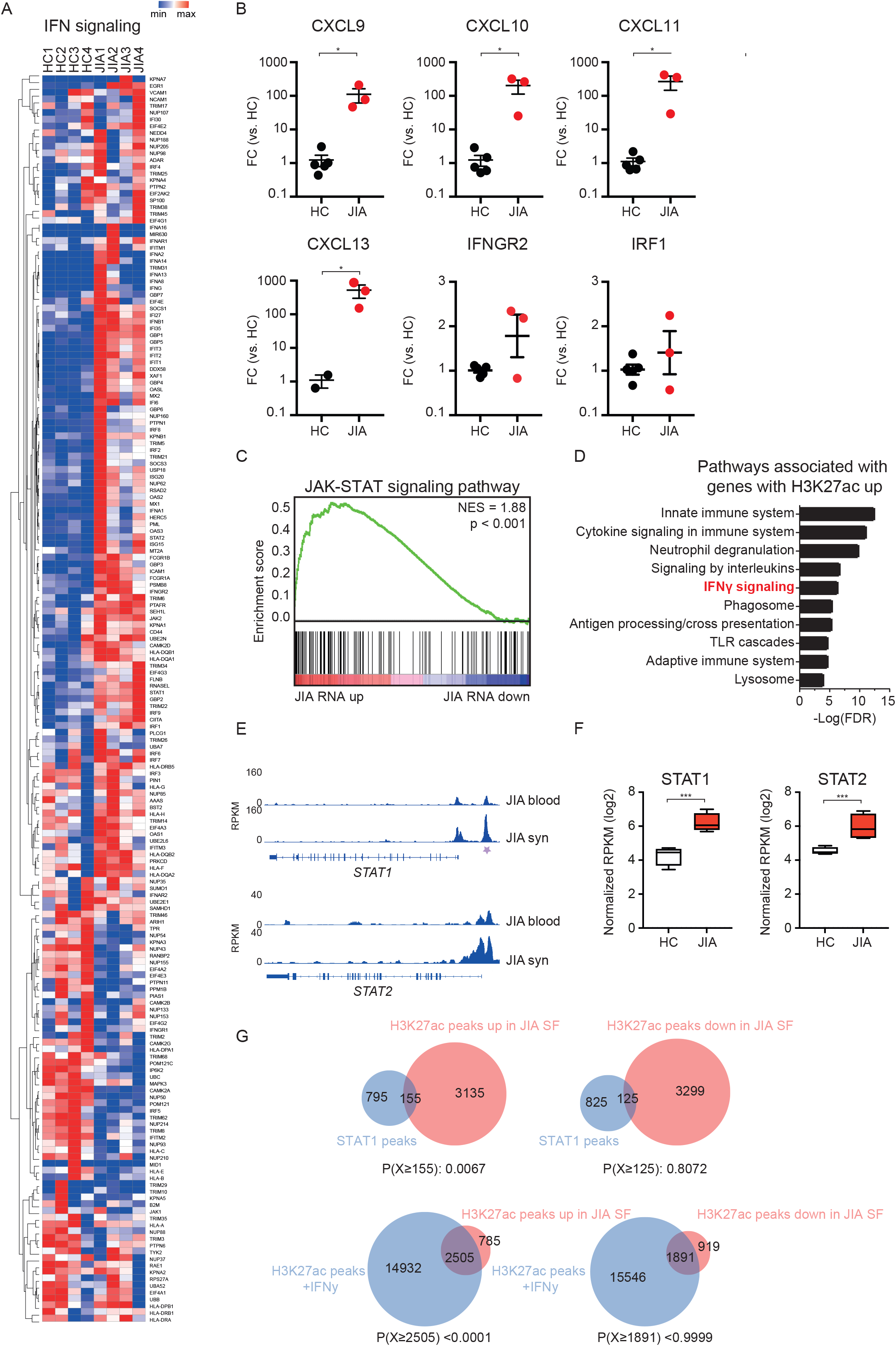
IFNγ-associated genes are increased in JIA SF monocytes. (**A**) Heatmap demonstrating the expression of IFNγ signaling-associated genes in HC PB and JIA SF monocytes. (**B**) Gene expression of selected IFNγ signaling-associated genes in HC PB monocytes and JIA SF monocytes. (**C**) Top 10 pathways associated with genes associated with an increased H3K27ac signal. (**D**) Gene set enrichment of JAK-STAT signaling pathway-associated genes and genes differentially expressed within JIA SF monocytes. (**E**) Gene track for *STAT1* and *STAT2* displaying H3K27ac signals for JIA PB and JIA SF. Purple asterisk indicates H3K27ac regions that are significantly different. (F) Normalized RPKM (log2) values of *STAT1* and *STAT2* expression in HC and JIA monocytes. (**G**) Venn diagrams displaying the overlap of H3K27ac peaks upregulated or downregulated in JIA SF monocytes with STAT1 or H3K27ac peaks of IFNγ-treated monocytes. STAT1 and H3K27ac data was obtained from GSE43036 (GSM1057010 and GSM1057016, respectively). P values for B were calculated using a Mann-Whitney test. P values for E were calculated using a hypergeometric test. * = p<0.05; ** = p<0.01, *** = p<0.001.

H3K27ac peaks that are increased in our inflammatory monocyte dataset were enriched for STAT1 binding and overlapped with H3K27ac signals mediated by IFNγ-treatment (Figure 3G). To experimentally assess the involvement of IFN and JAK-STAT signaling in establishing the epigenetic profile within inflammatory monocytes, we treated JIA SF monocytes for 4h with Ruxolitinib, an inhibitor of JAK1 and JAK2, and performed ChIP-seq for H3K27ac and H3K4me1 (*24*). While H3K27ac characterizes active enhancers, H3K4me1 is enriched at both active and inactive enhancer regions (*9*). This revealed that Ruxolitinib treatment induced approximately 450 significant changes in the H3K27ac epigenome, of which the majority of changes comprised a reduced H3K27ac signal (Figure 4A). Genes associated with a decrease in H3K27ac are involved in IFN signaling and included several components of the JAK-STAT signaling pathway, indicating an auto-regulatory feedback loop (Figure 4B and 4C). Ruxolitinib did not alter the H3K4me1 profile of inflammatory monocytes (Figure 4D). This is not surprising as histone methylation marks are less dynamically regulated than histone acetylation marks, which suggests that Ruxolitinib impairs the activity of active enhancers (*25*). Moreover, Ruxulitinib treatment preferentially decreased the signal of H3K27ac peaks that were increased in inflammatory monocytes (Figure 4E). Similarly, genes associated with a decreased H3K27ac signal upon Ruxolitinib treatment were expressed to a higher extent in inflammation-derived monocytes compared to HC monocytes (Figure 4F). Taken together, these data indicate that JAK inhibition preferentially reduces expression of genes associated with an increased H3K27ac signal in monocytes derived from the local inflammatory site.

**Figure 4:**
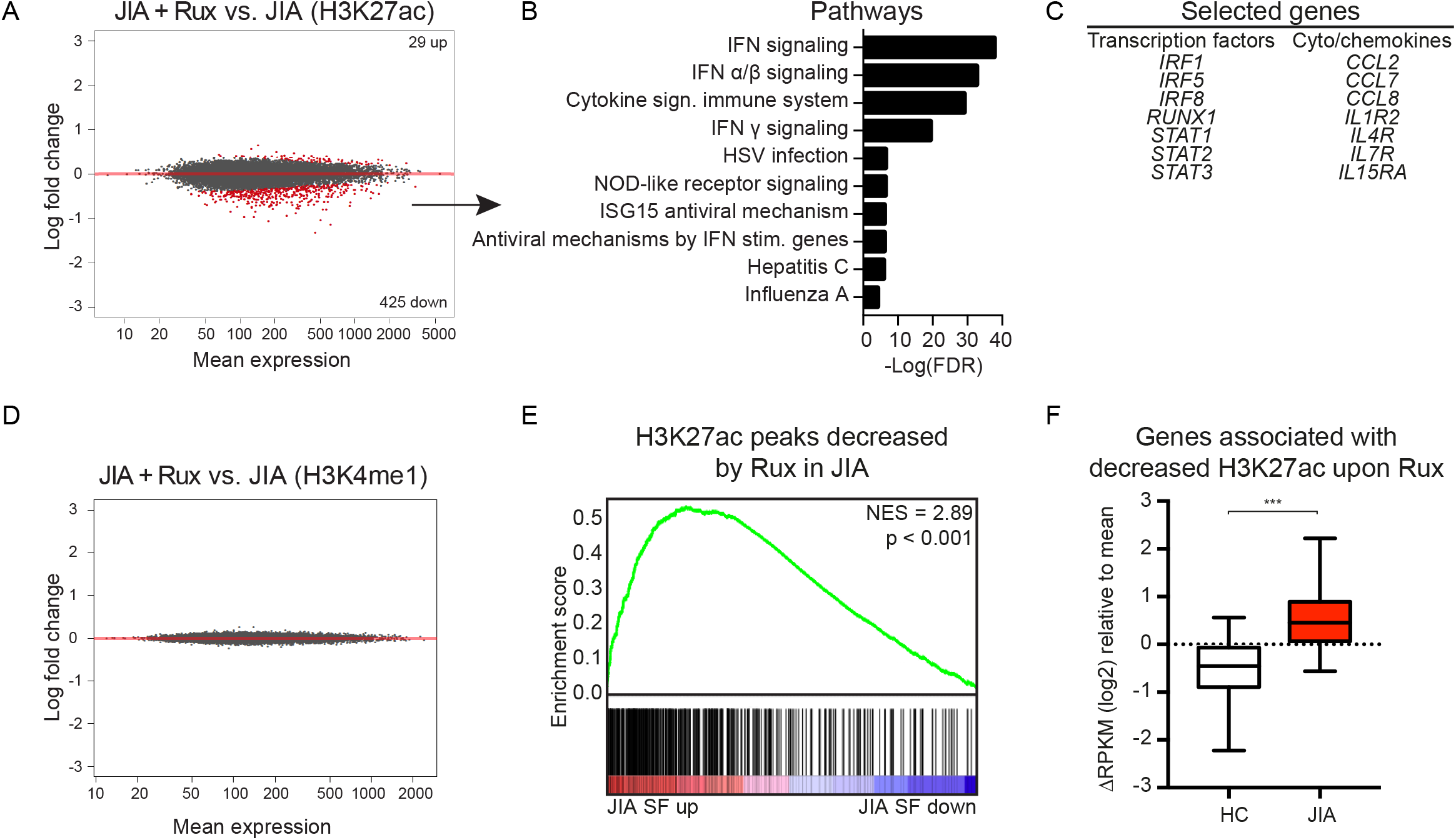
Ruxolitinib alters JIA SF monocytes at the epigenetic level. (**A**) MA plot of H3K27ac regions different within JIA SF monocytes after Ruxolitinib treatment. Red dot indicate genes with a FDR<0.1. (**B**) Top 10 pathways associated with genes significantly decreased by Ruxolitinib. (**C**) Selected genes significantly decreased by Ruxolitinib. (**D**) MA plot of H3K4me1 regions different within JIA SF monocytes after Ruxolitinib treatment. Red dot indicate genes with a FDR<0.1. (**E**) Gene set enrichment analysis of H3K27ac peaks decreased by Ruxolitinib in JIA and H3K27ac peaks different in JIA SF. (**F**) Boxplot with 5%-95% whiskers displaying ΔRPKM (log2) values of genes associated with a decreased H3K27ac signalin JIA monocytes after Ruxolitinib treatment. P value for E was calculated using a Wilcoxon-matched pairs signed rank test. * = p<0.05; ** = p<0.01, *** = p<0.001.

### Ruxolitinib inhibits the activated phenotype of JIA SF monocytes

Alteration of the epigenetic landscape by Ruxolitinib affected the activated phenotype of synovial monocytes, with the most predominant effect on IFN and cytokine signaling (Figure 5A, 5B, and Supplemental Figure 4A-4C). Genes decreased by Ruxolitinib are enriched within the genes that are upregulated in JIA, indicating that Ruxolitinib leads to reduced expression of disease-associated genes (Figure 5C and 5D). Furthermore, genes associated with an increased H3K27ac signal in inflammatory monocytes were preferentially decreased by Ruxolitinib, indicating that Ruxolitinib is targeting H3K27ac-regulated genes (Figure 5E). Expression of many cytokines and chemokines was increased in synovial monocytes compared to HC monocytes (Figure 1E). Ruxolitinib inhibits the expression of many of these cytokines, as demonstrated by RNA-sequencing and qPCR validation experiments (Figure 5F and 5G). To assess whether these observations could be extended to other local inflammatory site-derived monocytes, we assessed monocytes derived from the SF of RA patients. This revealed that RA-associated genes are also preferentially decreased by Ruxolitinib (Figure 5H), indicating that our observations are not specific for JIA but represent local autoinflammation as a whole. Taken together, these data demonstrate that Ruxolitinib-mediated inhibition of enhancer activity results in repression of the expression of genes associated with (auto-)inflammation in monocytes.

**Figure 5:**
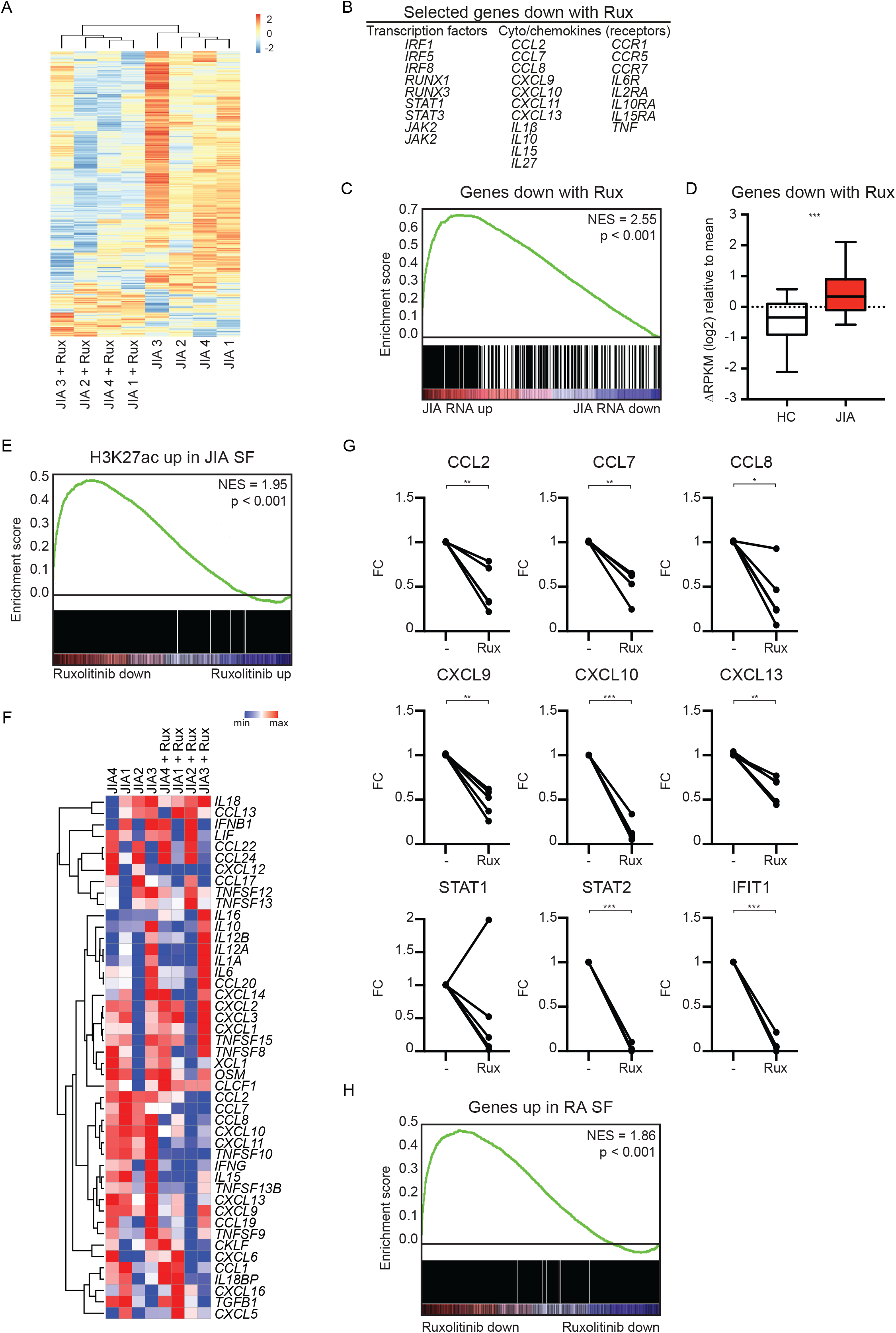
Ruxolitinib decreases the activated phenotype of JIA SF monocytes. (**A**) Supervised clustering analysis based on differences in gene expression in JIA SF monocytes after treatment with Ruxolitinib. (**B**) Selected genes downregulated by Ruxolitinib. (**C**) Gene set enrichment analysis of genes downregulated by Ruxolitinib and genes differentially expressed in JIA SF monocytes vs. HC monocytes. (**D**) Boxplot with 5%-95% whiskers displaying ΔRPKM (log2) values of genes decreased by Ruxolitinib in signal in HC and JIA monocytes. (**E**) Gene set enrichment analysis of genes associated with an increased H3K27ac signal in JIA SF monocytes and genes affected by Ruxolitinib treatment in JIA SF monocytes. (**F**) Heatmap demonstrating the expression of selected cytokines and chemokines in JIA SF monocytes with and without Ruxolitinib treatment, PB and JIA SF monocytes. (**G**) Gene expression of selected genes in JIA SF monocytes affected by Ruxolitinib treatment. P values for C and G were calculated using a Wilcoxon-matched pairs signed rank test. (**H**) Gene set enrichment analysis of genes increased in RA SF monocytes and genes affected by Ruxolitinib treatment in JIA SF monocytes. * = p<0.05; ** = p<0.01, *** = p<0.001.

## Discussion

Here, we assessed the transcriptional and H3K27ac profile of synovial monocytes isolated from the JIA patients. We observed that these inflammatory monocytes display an activated phenotype, illustrated by increased gene expression of surface activation markers and inflammatory cytokines and chemokines. This transcriptional phenotype was mirrored in the H3K27ac landscape, as expression of genes that are associated with an increased H3K27ac signal was increased. This suggests that epigenetic regulation contributes to disease-associated gene expression and thus disease pathogenesis.

For several autoimmune disease, such as RA and SLE, an increased frequency in intermediate monocytes (CD14^+^CD16^+^) has been described (*26, 27*). Furthermore, intermediate monocytes have been described to be increased in the PB and SF of enthesitis-related arthritis, a specific subset of JIA (*28*). The JIA patients used in this study do not have elevated levels of CD14^+^CD16^+^ in the PB, but the frequency of intermediate monocytes in the SF is significantly increased compared to HC and JIA PB. Therefore, we sorted CD14^+^CD16^+^ from JIA SF as well as HC PB and compared those for our transcriptional profiling. Unfortunately, this was not possible for the ChIP-sequencing experiments as there were insufficient numbers of intermediate monocytes from the PB of HC and JIA patients and therefore isolated CD14^+^ cells were used. Some of the epigenetic differences observed between inflammation-derived monocytes and PB monocytes might therefore be the result of a different ratio of classical (CD14^+^CD16^-^) monocytes vs. intermediate monocytes. However, since we did observe a strong correlation between the H3K27ac profile and gene expression of inflammatory monocytes, we assume that these differences do not drastically affect the epigenetic profile on a global level.

Our data demonstrated that JIA synovial monocytes are epigenetically distinct, regarding their H3K27ac status, from JIA PB and HC monocytes, while the latter are relatively similar. This raises the question whether the epigenetic alterations in inflammatory monocytes are cell-intrinsic or a reflection of the highly inflammatory synovial environment. IFNγ is one of the pro-inflammatory mediators present within the SF of oligoarticular, polyarticular as well as systemic JIA patients (*29*). Natural type I IFN producing cells have been detected in the synovial tissue of JIA patients, although IFNα levels in SF are reported to be low (*30*). There are some indications that IFNγ might play an important role within JIA pathogenesis. For example, increased responsiveness of PB-derived classical monocytes as well as naïve CD4^+^ T cells towards IFNγ stimulation, as indicated by increased phospho-STAT3 and pospho-STAT1 levels, respectively, has been described for polyarticular JIA (*31*). In addition, gene expression analysis of SFMCs of extended oligoarticular JIA patients revealed increased IFNγ levels compared to persistent oligoarticular JIA SFMCs, which corresponded with a trend of more IFNγ in the SF of these patients (*32*). Furthermore, IFNγ has been described to be produced by both CD4^+^ and CD8^+^ SF T cells and contributes to the resistance of SF CD8^+^ T cells to suppression (*33*). Our data indicated that transcriptional and epigenetic changes in inflammation-derived monocytes are associated with the IFN-signaling pathway. Together with the lack of H3K27ac differences between JIA PB and HC PB-derived monocytes, this suggests that monocytes are epigenetically altered at the site of inflammation due to cell-extrinsic factors, leading to altered gene expression which results in increased activation and cytokine production and thereby creating an auto-inflammatory loop. For our *in vitro* experiments using Ruxolitinib, we cultured synovial monocytes in the absence of other cell types or exogenous cytokines. Ruxolitinib-treatment affected the IFN signaling pathway, indicating that monocyte-derived IFN can activate monocytes itself and suggests that autocrine signaling contributes to the auto-inflammatory loop.

An epigenetic role for IFNγ and type I IFNs has been described in macrophages. IFNγ primes existing regulatory elements by increasing histone acetylation and chromatin remodeling, which leads to increased responsiveness of macrophages to pro-inflammatory stimuli, such as LPS or type I interferons (*23, 34*). This increased responsiveness is illustrated by increased transcription of *TNF* and *IL6*, pro-inflammatory cytokines which we found to be highly expressed by inflammatory monocytes. IFNα has a similar role and primes chromatin to enable robust transcriptional responses to weak signals (*35*). For example, IFNα enhances TNF inflammatory function by preventing the silencing of inflammatory genes. IFNγ can also induce *de novo* or latent enhancer formation by inducing transcription factors, such as STATs and Interferon Regulatory Factors (IRFs), that cooperate with lineage-determining transcription factors to open chromatin and deposit enhancer-associated histone marks in mouse macrophages (*36*). We observed that inhibiting IFN signaling in JIA synovial monocytes using the JAK inhibitor Ruxolitinib profoundly affected the H3K27ac landscape, while the H3K4me1 profile was unaffected. This indicates that in our experimental setting, Ruxolitinib predominantly affects existing regulatory elements by reducing H3K27ac deposition. This is in line with observations in T cells, where STATs have been demonstrated to have a major effect on enhancers bound by the histone acetyltransferase p300, while the impact of STATs on H3K4me1-positive enhancers was minimal (*37*). However, JAK inhibition prior to IFNγ stimulation has been demonstrated to affect H3K27ac as well as H3K4me1 at stimuli-specific enhancers (*36*). This suggests that, in patients, next to altering the established epigenetic landscape of monocytes at the site of inflammation, JAK inhibition might also affect formation of *de novo* enhancers. JAK inhibitors are currently being used for the treatment of RA, ulcerative colitis, and psoriasis and recently the JAK inhibitor Tofacitinib has been approved for the treatment or patients with active polyarticular course JIA (*38, 39*). The latter is based on a phase 3 study evaluating the efficacy and safety of Tofacitinib which demonstrated that the occurrence of disease flares was significantly decreased in Tofacitinib-treated patients compared to placebo-treated patients (*40*–*42*). Our data indicate that the therapeutic effect of JAK inhibition in JIA is, at least partly, established by altering the epigenetic landscape of inflammatory monocytes, thereby reducing the expression of disease-associated genes. Gene set enrichment analysis demonstrated that genes upregulated in RA synovial-derived monocytes are enriched within the genes downregulated by Ruxolitinib in JIA, indicating that these genes in RA might also be JAK-dependent and regulated by H3K27ac (*43*). It will be interesting to study if JAK inhibitors have a similar effect in other autoimmune diseases as well. Furthermore, the development of next-generation JAK-inhibitors that selectively inhibit JAK2 will allow for more precise control of disease-associated gene expression, as JAK2 is upregulated in inflammatory monocytes (*39*). Altogether our data provide further rationale for the treatment of autoimmune disease patients with JAK inhibitors and indicate that other molecules that affect the epigenetic landscape of inflammatory monocytes, or other immune cells, can have a therapeutic effect as well.

## Materials and methods

### Collection of synovial fluid (SF) and peripheral blood (PB) samples

Sixteen oligoarticular JIA patients were included in this study who at the time of sampling all had active disease. PB and SF samples were obtained at the same moment, either via vein puncture or intravenous drip and therapeutic joint aspiration, respectively. Informed consent was obtained from all patients either directly or from parents/guardians when the patients were younger than age 12 years. The study procedures were approved by the Institutional Review Board of the University Medical Center Utrecht (UMCU; METC nr: 11-499/C) and performed according to the principles expressed in the Helsinki Declaration. SF samples were treated with hyaluronidase (Sigma-Aldrich) for 30 min, 37°C. Synovial fluid mononuclear cells (SFMCs) and peripheral blood mononuclear cells (PBMCs) were isolated using Ficoll-Paque density gradient centrifugation (GE Healthcare) and were either used immediately or frozen and stored at -80°C until further use.

### Monocyte isolation and culture

For RNA-sequencing of *ex vivo* HC and JA monocytes, HC PBMCs and JIA SFMCs were thawed, CD3^+^ cells were depleted by human anti-CD3 microbeads (Milteny Biotec) according to the manufacturer’s instructions and CD14^+^CD16^+^ cells were sorted by flow cytometry using a BD FACS Aria III (BD Biosciences). For validation of *ex vivo* sequencing results, CD14+ were isolated from frozen HC PBMCs and JIA SFMCs by magnetic-activated cell sorting (MACS) using human CD14 microbeads (Milteny Biotec) according to the manufacturer’s instructions. For ChIP-sequencing of *ex vivo* HC and JA monocytes (Figure 2), HC PBMCs, JIA PBMCs and JIA SFMCs CD14^+^ cells where isolated by flow cytometry using a BD FACS Aria II (BD Biosciences). For analysis of JQ1 sensitive genes, HC PBMCs were thawed and CD14^+^ cells were sorted by flow cytometry using a BD FACS Aria II (BD Biosciences). Subsequently, CD14^+^ cells were cultured o/n with 100 ng/mL LPS (Invivogen) and treated with 300 nM JQ1(-) or JQ1(+) (ApexBio) in RPMI 1640 + GlutaMAX (Life Technologies) supplemented with 100 U/ml penicillin (Gibco), 100 mg/ml streptomycin (Gibco) and 10% heat-inactivated human AB-positive serum (Invitrogen) at 37°C in 5% CO_2_. For RNA-and ChIP-sequencing of Ruxolitinib-treated samples, CD14^+^ cells were isolated by magnetic-activated cell sorting (MACS) from JIA SFMCs and cultured for 4h with or without 1 μM Ruxolitinib (Selleck Chemicals).

### ChIP-sequencing and analysis

Cells were crosslinked with 2% formaldehyde (Sigma-Aldrich) and after 10 min crosslinking was stopped by adding 0.15 M glycine. Nuclei were isolated in in 50 mM Tris (pH 7.5), 150 mM NaCl, 5 mM EDTA, 0.5% NP-40, and 1% Triton X-100 and lysed in 20 mM Tris (pH 7.5), 150 mM NaCl, 2 mM EDTA, 1% NP-40 and 0.3% SDS. Lysates were sheared using Covaris microTUBE (duty factor 20%, peak incident power 105, 200 cycles per burst, 480 sec cycle time) and diluted in 20 mM Tris (pH 8.0), 150 mM NaCl, 2 mM EDTA, 1% Triton X-100. Sheared DNA was incubated overnight with anti-histone H3 acetyl K27 antibody (ab4729; Abcam) pre-coupled to protein A/G magnetic beads (Pierce). Samples were washed and crosslinking was reversed by adding 1% SDS, 100 mM NaHCO3, 200 mM NaCl, and 300 ug/ml proteinase K (Invitrogen). DNA was purified using ChIP DNA Clean & Concentrator kit (Zymo Research), endrepair, a-tailing, and ligation of sequence adaptors was done using Truseq nano DNA sample preparation kit (Illumina). Samples were PCR amplified and barcoded libraries were sequenced 75 bp single-end on Illumina NextSeq500. Peaks were called using Cisgenome 2.0 (–e 150 -maxgap 200 –minlen 200) (*44*). Peak coordinates were stretched to at least 2000 base pairs and collapsed into a single list. Overlapping peaks were merged based on their outmost coordinates. Only peaks identified by at least 2 independent datasets were further analyzed. Peaks with differential H3K27ac occupancy were identified using DESeq (padj<0.1) (*45*). Associated genes were defined as genes with a transcription start site within 20 kB from the H3K27ac peak. To assess enrichment of STAT1 and H3K27ac ChIP-seq data of IFNγ-treated monocytes (GSE43036) in JIA SF H3K27ac peaks, peak coordinates of GSM1057010 and GSM1057016 were overlapped with significantly upregulated or downregulated genes in JIA SF vs. JIA Blood.

### RNA-sequencing and analysis

For analysis of JQ1 sensitive genes, RNA-sequencing was performed as described before (Cell Reports). For analysis of *ex vivo* gene expression, total RNA was extracted using Trizoll, mRNA was isolated using Poly(A)Purist MAG kit (Life Technologies) and additionally purified with a mRNA-ONLY Eukaryotic mRNA Isolation Kit (Epicentre). Transcriptome libraries were then constructed using SOLiD total RNA-seq kit (Applied Biosystems) and sequenced using 5500 W Series Genetic Analyzer (Applied Biosystems) to produce 40-bp-long reads. Reads were aligned to the human reference genome GRCh37 using STAR version 2.4.2a. Picard’s AddOrReplaceReadGroups (v1.98) was used to add read groups to the BAM files, which were sorted with Sambamba v0.4.5 and transcript abundances were quantified with HTSeq-count version 0.6.1p1 using the union mode. Subsequently, reads per kilobase million reads sequenced (RPKMs) were calculated with edgeR’s RPKM function. Differentially expressed genes were identified using the DESeq2 package with standard settings. Genes with padj<0.05, with a base mean ≥ 10 and a log2 FC ≥1 or ≤ -1 were considered as differentially expressed and used for further analysis.

For analysis of Ruxolitinib-affected gene expression, total RNA was purified using the PicoPure RNA Isolation kit (Thermo Fisher Scientific). mRNA was isolated using NEXTflex^®^Poly(A) Beads (Bio Scientific), libraries were prepared using the NEXTflex^®^Rapid Directional RNA-Seq Kit (Bio Scientific) and samples were sequenced 75 base pair single-end on Illumina NextSeq500 (Illumina, Utrecht DNA Sequencing Facility) and processed as described above. Genes with absolute padj<0.1 were considered as differentially expressed and used for further analysis.

### Gene ontology enrichment analysis and gene set enrichment analysis (GSEA)

Gene ontology (GO) enrichment analysis was performed using the ToppFunn tool of ToppGene Suite(*46*). GO terms with an adjusted (Benjamini-Hochberg procedure) p-value <0.05 were considered being enriched. Gene set enrichment analysis was performed using GSEA pre-ranked(*47*). Significance of the enrichment was calculated based on 1000 cycles of permutations and the normalized enrichment score (NES) and the p-value (FDR) are annotated.

### Quantitative RT-PCR

Total RNA was extracted using the PicoPure RNA Isolation kit and cDNA synthesis was performed using the iScript cDNA synthesis kit (Bio-Rad). cDNA samples were amplified with SYBR Select mastermix (Life Technologies) in a QuantStudio 12k flex (Thermo Fisher Scientific) according to the manufacturer’s protocol.

### Multiplex immunoassay

Multiplex analysis (xMAP; Luminex) was performed as described previously on supernatant derived from HC and JIA-derived CD14+ cells either cultured o/n (to determined *ex vivo* cytokine production) or for 4h in the presence of 1 μM Ruxolitinib (de Jager et al., 2005).

### Data availability

The RNA-sequencing and ChIP-sequencing data from this publication have been deposited in the NCBI GEO database and together assigned the identifier GSEXXXX.

## Figure legends

**Supplemental Figure 1: JIA SF monocytes are transcriptionally different**. (**A**) Gene expression of selected surface markers in JIA SF monocytes vs. HC PB monocytes. P values for B were calculated using a Mann-Whitney test. * = p<0.05; ** = p<0.01, *** = p<0.001.

**Supplemental Figure 2: JIA SF monocytes are epigenetically different**. (**A**) MA plot of genes differentially expressed between JIA PB and HC PB monocytes. Red dots indicate genes that are significantly different. (**B**) MA plot for genes differentially expressed in JIA SF monocytes after JQ1 treatment. Red dots indicate genes that are significantly different. (**C**) Top 10 biological processes associated with genes that are significantly decreased by JQ1 treatment.

**Supplemental Figure 3: Expression levels of IFN**γ**-induced chemokines**. (**A**) Protein levels of selected chemokines in JIA SF monocytes and HC PB monocytes. P values were calculated using a Mann-Whitney test. * = p<0.05; ** = p<0.01, *** = p<0.001.

**Supplemental Figure 4: Ruxolitinib decreases the activated phenotype of JIA SF monocytes**. (**A**) MA plot of genes differentially expressed within JIA SF monocytes after Ruxolitinib treatment. Red dot indicate genes with a FDR<0.1. (**B**) Top 10 pathways associated with genes downregulated by Ruxolitinib in JIA SF monocytes. (**C**) Top 3 diseases associated with genes downregulated by Ruxolitinib in JIA SF monocytes.

