## Supplementary figures for "Epigenetic changes in autoimmune monocytes contribute to disease and can be targeted by JAK inhibition"

A

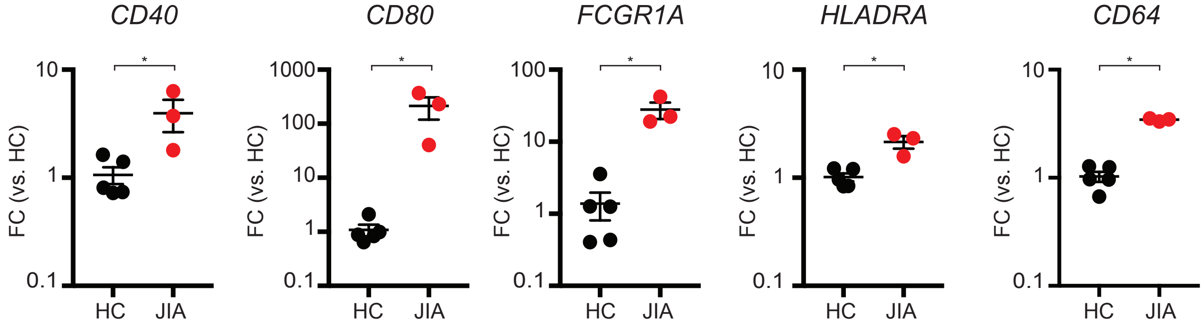
CD40
CD80
FCGR1A
HLADRA
CD64
FC (vs. HC)
10
1
0.1
1000
100
10
1
0.1
*
HC
JIA

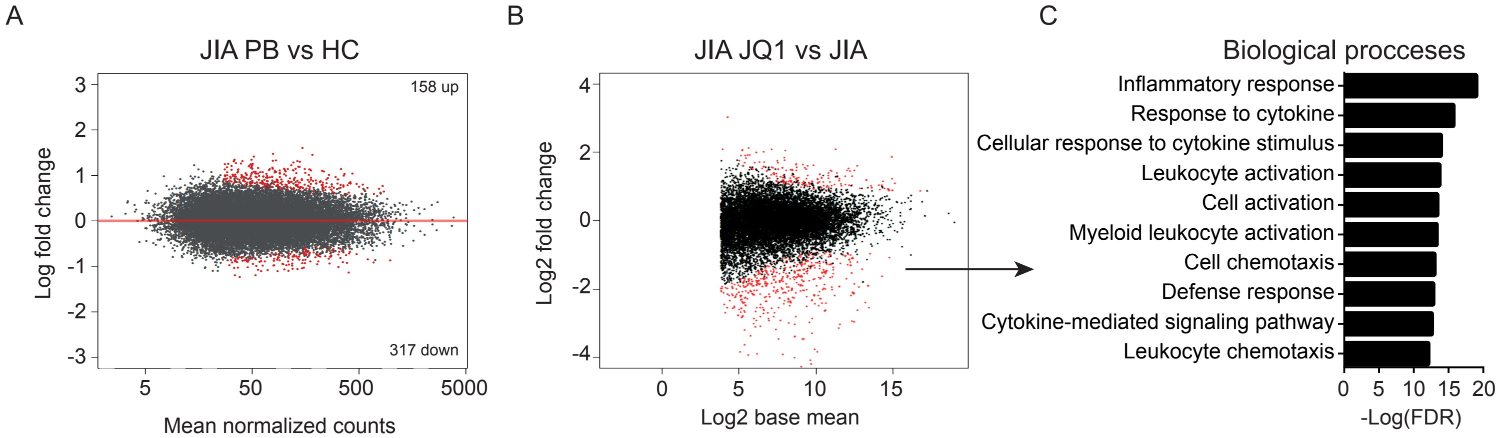


No enrichment of biological processes

Supplemental Figure 2 | Peeters *et al.,* 2021

A

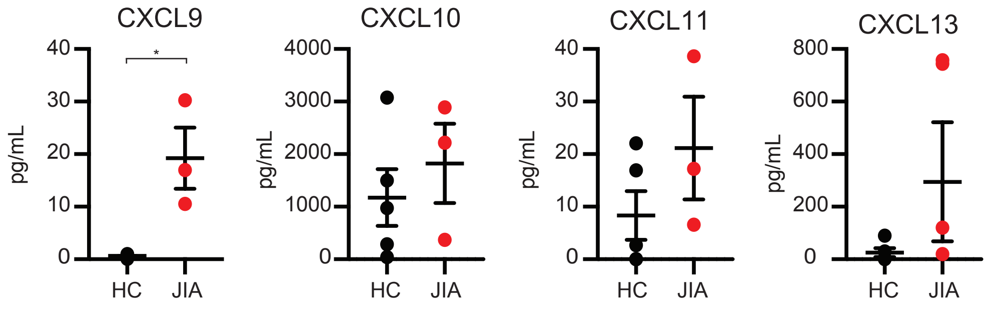


Supplemental Figure 3 | Peeters *et al.,* 2021

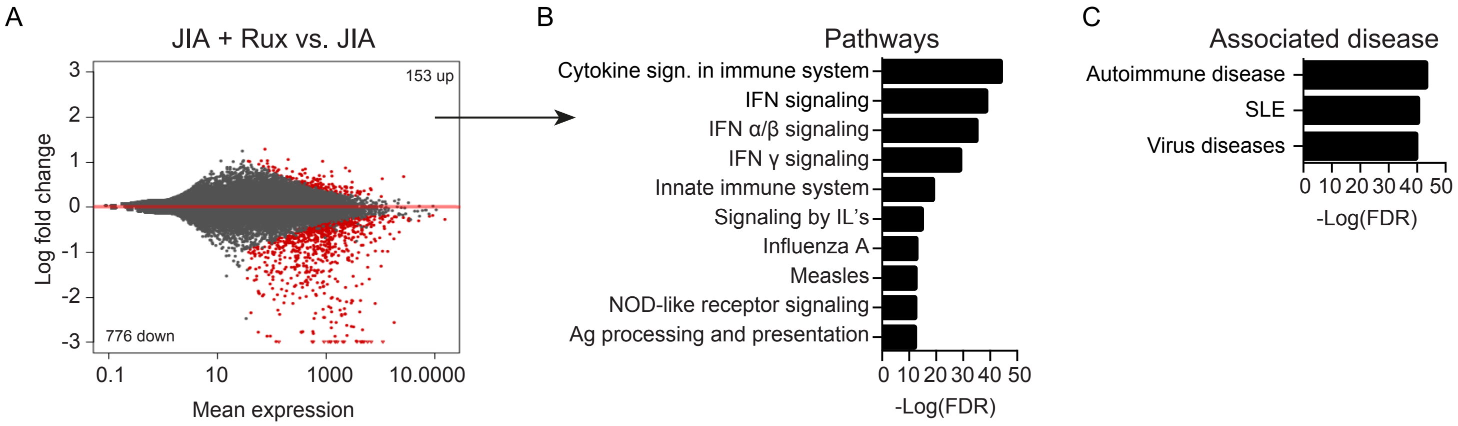


Supplemental Figure 4 | Peeters *et al.,* 2021
